# A nanometer difference in myofilament lattice spacing of two cockroach leg muscles explains their different functions

**DOI:** 10.1101/656272

**Authors:** Travis Carver Tune, Weikang Ma, Thomas C. Irving, Simon Sponberg

## Abstract

Muscle is highly organized across scales. Consequently, small changes in arrangement of myofilaments can influence macroscopic function. Two leg muscles of a cockroach, have identical innervation, mass, twitch responses, length-tension curves, and force-velocity relationships. However, during running, one muscle is dissipative, while the other produces significant positive mechanical work. Using time resolved x-ray diffraction in intact, contracting muscle, we simultaneously measured the myofilament lattice spacing, packing structure, and macroscopic force production of these muscle to test if nanoscale differences could account for this conundrum. While the packing patterns are the same, one muscle has 1 nm smaller lattice spacing at rest. Under isometric activation, the difference in lattice spacing disappeared explaining the two muscles’ identical steady state behavior. During periodic contractions, one muscle undergoes a 1 nm greater change in lattice spacing, which correlates with force. This is the first identified feature that can account for the muscles’ different functions.

## Introduction

Many biological structures, especially tissues have hierarchical, multiscale organization (***McCulloch, 2016***). Of these, muscle is exceptional because it is also active: capable of producing internal stress based on the collective action of billions of myosin motors (***Maughan and Vigoreaux, 1999***). Muscle can perform many roles in organisms, acting like a motor, brake, or spring depending on the task required (***Josephson, 1985***; ***Dickinson et al., 2000***). It is even possible for different parts of a single muscle to behave with different functions (***Roberts et al., 1997***; ***George et al., 2013***). This energetic versatility enables muscle’s diverse function in animal locomotion and behavior. Yet we still have a difficult time predicting function from multiscale properties.

Muscle function during locomotion is typically characterized through a work loop: a stress-strain (or force-length) curve in which the length (or strain) of the muscle is prescribed through a trajectory and electrically activated at specific points (phases) during the cycle of shortening and lengthening (***Josephson, 1985***; ***Ahn, 2012***). The area inside the loop gives the net work done by the muscle and can be positive, negative, biphasic, or zero. Work loop parameters typically mimic *in vivo* or power maximizing conditions. Many physiological characterizations of muscle are steady state in some respect – twitch responses are isometric, the length-tension curve is obtained under constant, usually tetanic activation, and even the force-velocity curve is taken as the force at constant activation during constant velocity shortening for a given load. These macroscopic properties arise from and, in fact, helped establish the crossbridge basis for muscle contraction and sliding filament theory (***Gordon et al., 1966***; ***Huxley and Simmons, 1971***). Although these steady state macroscopic measurements are important determinants of muscle work loops, they are not sufficient to account for the variability of muscle work output and hence function under dynamic conditions (***Josephson, 1999***). The multiscale nature of muscle suggests that subtle differences in structure of the contractile apparatus at the micro to nanometer scale could also be playing an underappreciated role in determining differences in work output and hence macroscopic function.

Differences at the nanometer scale can have profound effects due to the arrangement of actin-containing thin filaments and myosin-containing thick filaments into a regular lattice with spacings on the scale of 10’s of nanometers (***Millman, 1998***). This myofilament lattice inside each sarcomere is a crystal in cross section even under physiological conditions. As a result, its structure can be readily studied by x-ray diffraction even during force production and length changes (***Irving, 2006***; ***Iwamoto, 2018***). The interfilament spacing within the lattice (lattice spacing) depends in part on the axial length of the muscle, stemming from the strain placed on the muscle fibers during contraction, as well as the radial tension (***Bagni et al., 1994***). Lattice spacing in turn is important for force development because it influences myosin binding probability and hence axial and radial force production (***Schoenberg, 1980***; ***Williams et al., 2010***; ***Tanner et al., 2007, 2012***). Lattice spacing changes in muscle independent of sarcomere length changes have been shown to enhance Ca^2+^ sensitivity (shape of force-pCa curves) (***Fuchs and Wang, 1996***) and change crossbridge kinetics ***Adhikari et al. (2004)***. The change in lattice spacing even accounts for up to 50% of the force change due to length in a typical muscle’s force-length curve (***Williams et al., 2013***). The filament lattice in muscle is not isovolumetric, indicating crossbridge attachment generates a radial force which corresponds to and is of the same order of magnitude as crossbridge axial force (***Bagni et al., 1994***; ***Cecchi et al., 1990***). These studies all showed how lattice spacing could affect macroscopic properties of muscle, but the implications have so far only examined steady state or quasi-static conditions. However, significant variation in lattice spacing has been linked to crossbridge binding during work loops in isolated insect flight muscle where temperature was changed to affect both lattice spacing and work (***George et al., 2013***). What is still unknown is whether or not myofilament lattice structure (its packing arrangement and spacing) is a significant determinant of macroscopic work in the absence of other effects, and if differences in lattice structure result in a difference in muscle work in a manner functional for locomotion.

To explore these questions we looked for two very similar muscles that have unexplained differences in their work production. Two of the femoral extensors of the cockroach, *Blaberus discoidalis*, are ideal for exploring how multiscale mechanisms influence work (Figure 1a). These two muscles have the same submaximal and tetanic force-length curves, twitch response, force-velocity curve, phase of activation, force enhancement due to passive pre-stretch, and force depression due to active shortening (***Full et al., 1998***; ***Ahn et al., 2006***). They are even innervated by the same single, fast-type motor neuron (***Becht and Dresden, 1956***; ***Pearson and Iles, 1971***) and share the same synaptic transmission properties (***Becht et al., 1960***) meaning that both muscles are activated as a single motor unit in all conditions. To the best of our knowledge, these muscles share the same anatomical and steady state physiological properties. However, when the muscles are isolated and prescribed dynamic patterns of strain and activation which match those that the muscle experiences during *in vivo* running, one muscle acts like a brake with a dissipative work loop, while the other is more like a motor with a net positive, biphasic work loop (Figure 1b, work loops from ***Ahn et al. (2006)***). Since the macroscopic properties that might determine muscle function are the same in these muscles, we cannot account for their differences in work output. It has been suggested, although not tested, that structural differences in the myofilament lattice may account for the differences (***Ahn et al., 2006***).

**Figure 1.**
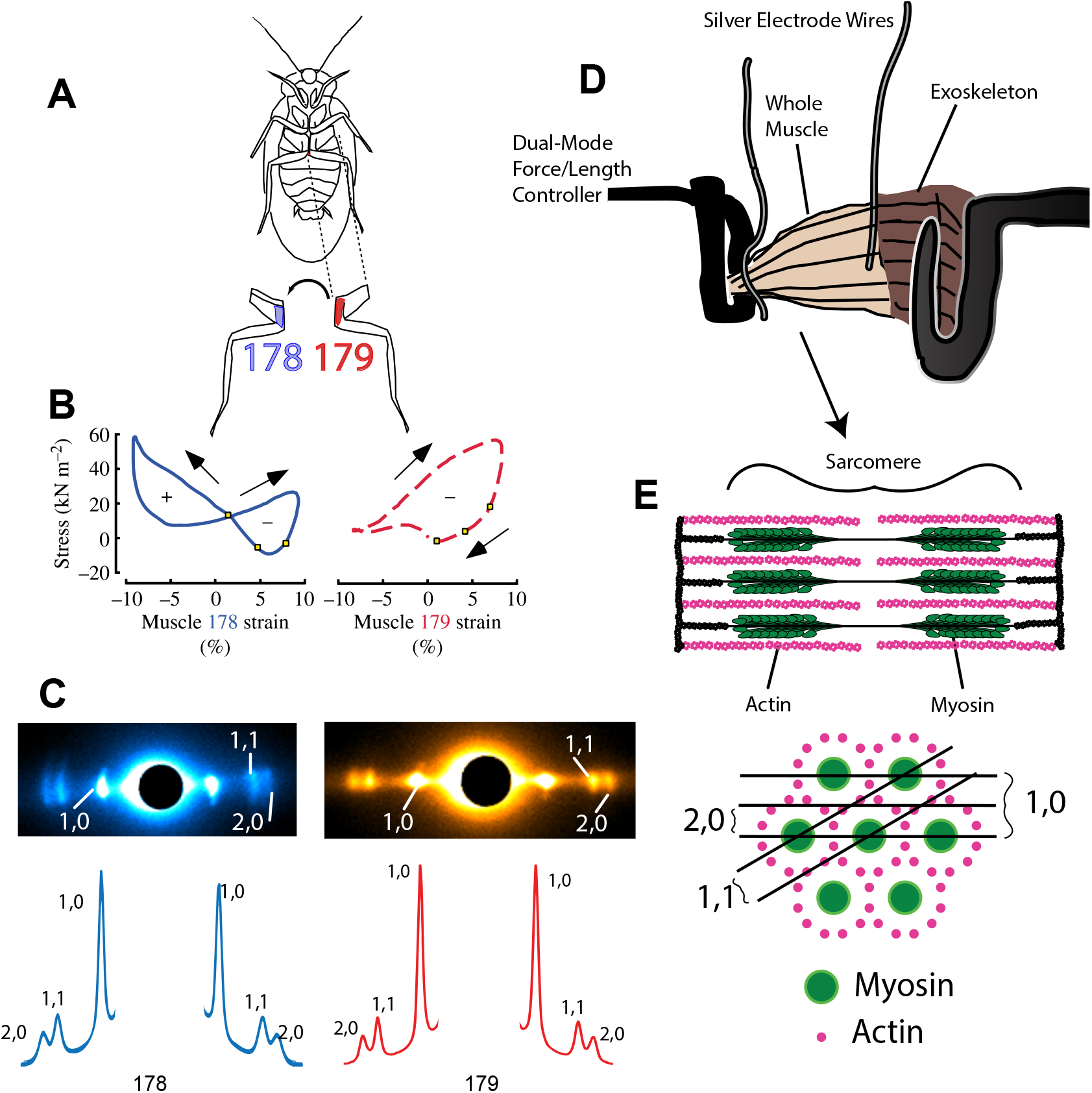
A) Ventral View of *Blaberus discoidalis* showing the hind-limb femoral extensors 178 and 179 (notation from ***Carbonell (1947)***). B) Work loops performed on muscles 178 and 179 show a difference in function despite near identical steady state behavior (work loop figures from ***Ahn et al. (2006)***). C) X-ray diffraction patterns from muscles 178 and 179 with the most prominent peaks labeled. Also shown, is the intensity profile along the equatorial axis. D) Diagram showing experimental set-up. X-ray beam path is perpendicular to the contraction axis. E) Multiscale hierarchy of muscle structure, showing a single sarcomere (1-10 *μ*m) of a muscle (1-10 mm) and the sarcomere cross-section, with diffraction planes (10’s of nm) corresponding to the peaks indicated in C. Spacing between diffraction planes in E is related by Bragg’s Law to the spacing between peaks in C, while the intensity of peaks shown in C are related to the mass lying along depicted planes in E.

Critically any feature than could explain the differences in work output would not only have to explain the dynamic differences between the two muscles, but must also be identical in steady state in both muscles in order to account for their similarities. We explore two hypotheses using time-resolved x-ray diffraction measurements of muscle’s nanometer structure and myofilament lattice spacing (Figure 1C) taken simultaneous with physiological force measurements in intact, contracting muscle (Figure 1D). First, we tested whether the lattice packing structure of the two muscles might be different. Actin and myosin vary in their ratio and the phase of their packing pattern across muscles (***Millman, 1998***; ***Squire et al., 2005***). Different packing structures could produce different dynamics of force development by affecting myosin free energy. Second, we consider if the myofilament lattice spacing (Figure 1E) is systematically different in the two muscles thereby affecting work production. If any structural differences only exists under dynamic conditions (periodic contractions), then they could also lead to convergent steady state properties.

## Results

### Similarity in packing structure cannot explain functional differences

We first tested whether the two muscles had the same lattice packing structure (Figure 1E). In invertebrates, there can be a wide variety of actin packing patterns. Two muscles with different myosin-actin ratios and geometry might have similar steady state behavior since they have the same number of myosin heads available for crossbridge binding, but could have different dynamic behavior because of differences in actin availability. We can use the ratio 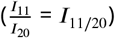 of intensity in the (1,1) and (2,0) peaks (Figure 1, peaks labeled) to determine if muscles 178 and 179 have similar packing patterns (see methods).

We measured the intensity of the (1,1) and (2,0) peaks of muscles 178 and 179 and found *I*_11/20_ = 2.47 ± 0.4 and *I*_11/20_ = 2.68 ± 0.4 for muscle 179 (mean and 95% confidence of mean) for muscles 178 and 179 respectively. We know from previous electron microscopy work that muscle 137, the midlimb analog of 179, has a 6:1 packing pattern common among insect limb muscle (***Jahromi and Atwood, 1969***), so it is likely muscle 179 also has this packing pattern. Regardless, based on the intensity ratio of 178 compared to 179, we determined 178 to have the same structure as 179. Since the two muscles have the same packing structure, this alone cannot account for their different work loops.

### A 1 nm difference in lattice spacing under passive conditions disappears when muscles are activated to steady state

Since we did not observe a difference in packing structure between the two muscles, we next asked if the lattice spacing under isometric conditions differed between the two muscles. The distance between myosin planes is proportional to the lattice spacing *d*_10_, which we can 1nd by measuring the distance between the corresponding diffraction peaks, *s*_10_, and using Bragg’s Law, 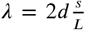, where L is the sample to detector distance and *λ* is the wavelength of the x-ray. At each strain condition, we isometrically held the muscle and activated with a 3 spike stimulus, reflecting the 3 spikes typical of submaximal activation in these muscles (***Ahn et al., 2006***). We used the value of *d*_10_ at peak stress as the steady state active *d*_10_.

At rest, passive 178 and 179 lattice spacings were different with 178 being 1.01 ± 0.41 nm (mean ± 95% CI of the mean) smaller across all 5 strain conditions (*p* = .005). When activated the myofilament lattice of muscle 178 expanded radially by about 1 nm across the entire strain range measured between passive and active conditions, while in 179 activation caused no statistically significant change in lattice spacing under any strain condition (Figure 3, *p* = 0.008 and *p* = 0.52, two-factor ANOVA, for 178 and 179 respectively). We also found that activated 178 and 179 lattice spacings were only .05 nm ± .4 apart (mean ± 95% CI of the mean) and were not significantly different (*p* = 0.86). Taken together, these measurements show a statistically significant difference between passive muscle 178 and 179, which disappears upon activation as 178’s *d*_10_ increases to match 179’s *d*_10_ which does not change.

**Figure 2.**
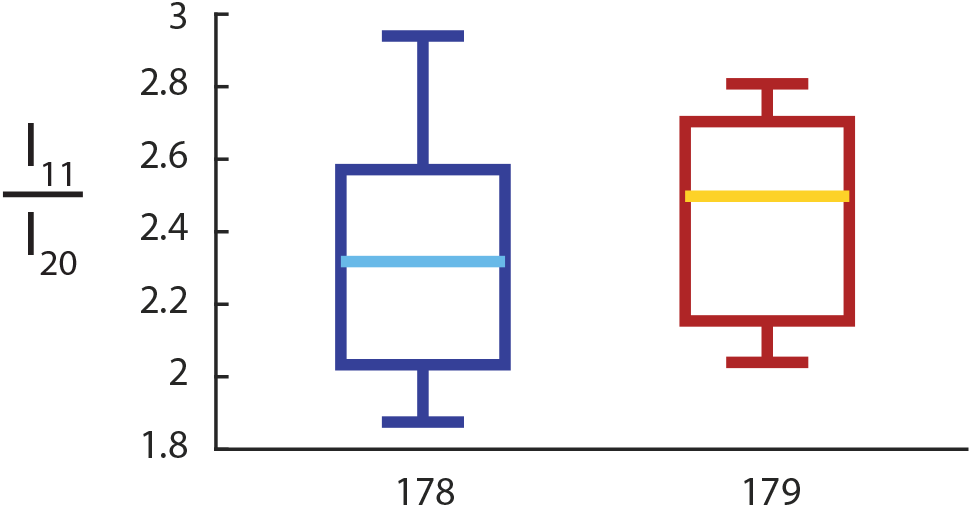
Boxplots of the intensity ratio *I*_11/20_ for muscles 178 (n=8, left) and 179 (n=9, right), with median and 25^th^ and 75^th^ percentiles. There is no significant difference between the two muscles’ intensity ratios, indicating that they have same packing pattern (*p* =.44, Wilcoxon rank sum test).

**Figure 3.**
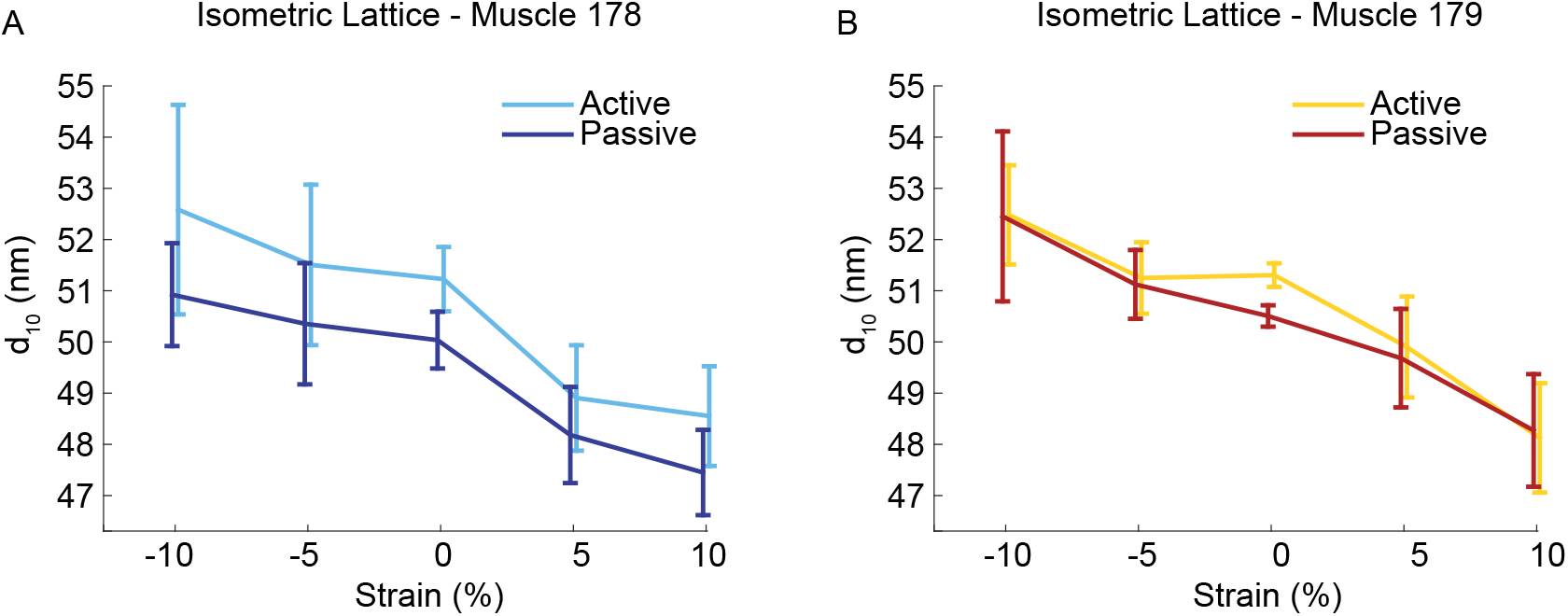
Muscle 178 (A) and 179 (B) passive and active *d*_10_ at strains of −10% to +10% of operating length, with 95% confidence of the mean. Sample size n at strains (−10,−5,0,5,10) was: (7,6,8,7,7) for muscle 178; (8,9,8,9,9) for muscle 179

### The two muscles have different lattice spacing dynamics

While the isometric differences are informative concerning potential differences in stress development, we also needed to examine how lattice spacing behaves during dynamic contractions. We tested conditions similar to the those where *in vivo* work is being generated to compare to ***Ahn et al. (2006)***. We measured *d*_10_ during passive work loops and work loops with the *in vivo* activation pattern and phase (see methods).

When activated, the time course of *d*_10_ in muscle 178 differed significantly in the active vs. the passive case, while 179 lattice spacing did not (*p* = .008 and *p* = .11, two factor ANOVA, see Figure 4). In both muscles passive (unstimulated) muscle underwent comparable lattice spacing change. Activation produced additional lattice spacing expansion of 1.1 ± .5 nm at the peak stress plateau. Peak lattice spacing change in muscle 179 was .4 ± .4 nm (see Figure 5 for a representative lattice spacing, stress, and incremental work timeseries).

**Figure 4.**
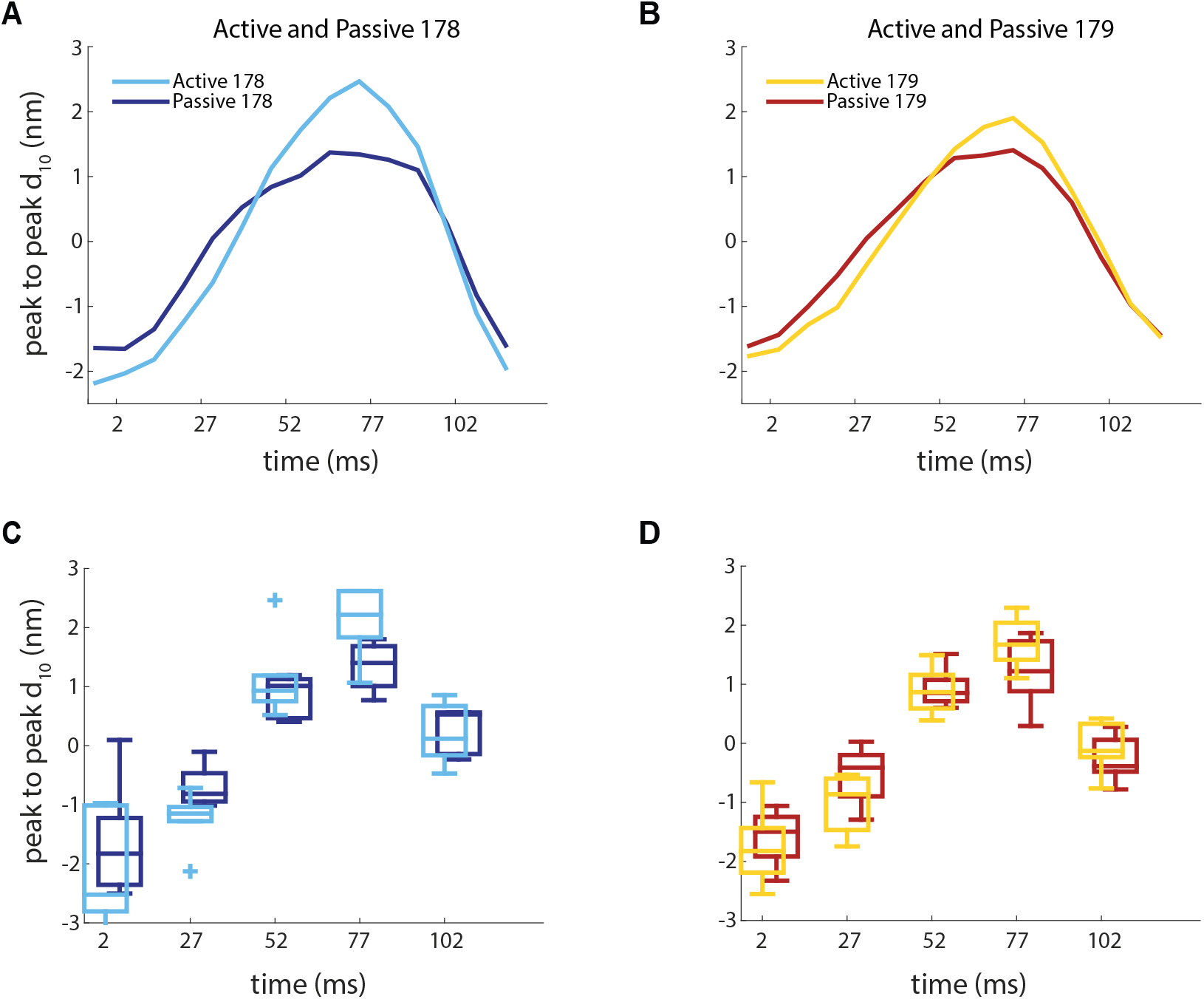
A) and B) show the mean subtracted active and passive *d*_10_ lattice spacing, respectively. These were obtained similarly to Figure 3, but under dynamic work loop conditions. C) and D) show the variation in the mean at times corresponding to .02*T,* 0.23*T,* 0.43*T,* 0.64*T,* 0.84*T,* which corresponded to the time points nearest maximum strain amplitude 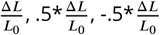, minimum strain amplitude 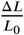, and 0% strain, respectively, where T = 120 ms is the cycle period. Boxplots show the median spacing as well as 25^th^ and 75^th^ percentiles. Sample size n was: 5 for muscle 178 passive, 6 for muscle 178 active, 8 for muscle 179 active and passive.

**Figure 5.**
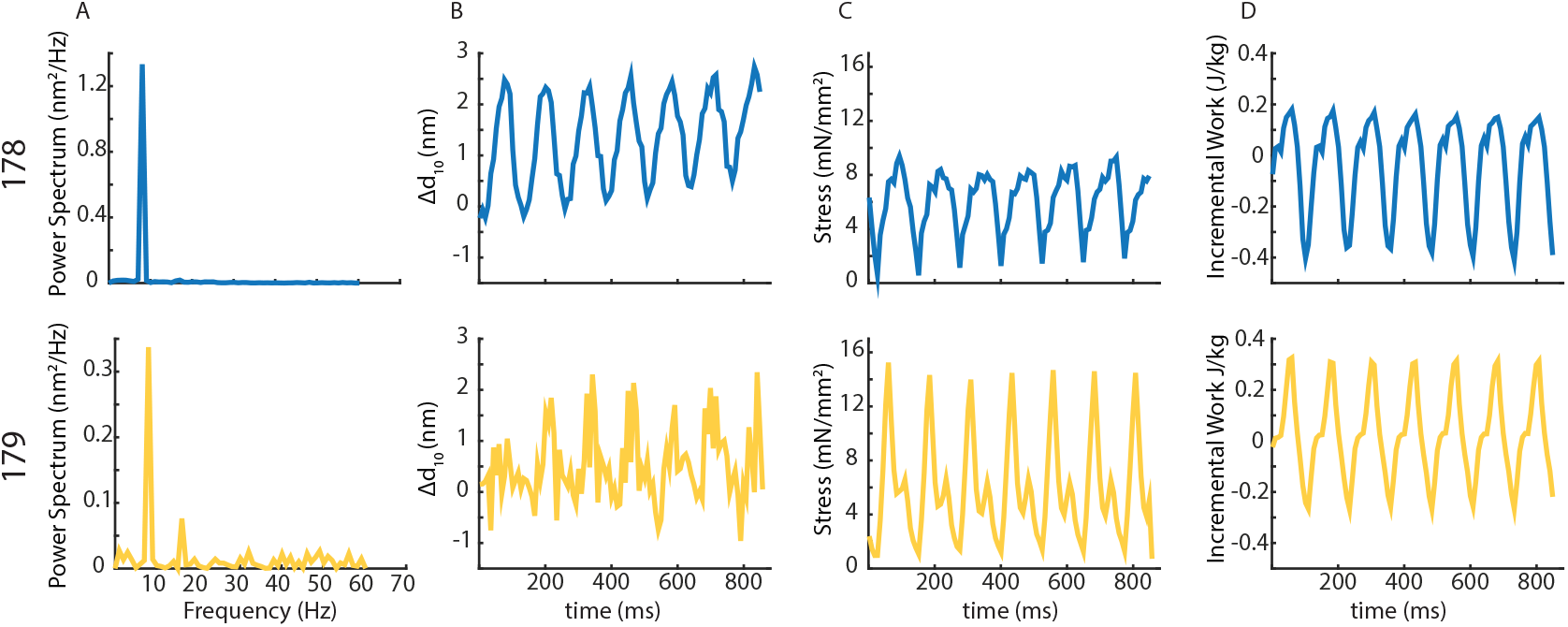
A) We calculated the Fourier series for Δ*d*_10_ for each trial and found significant power in the work loop frequency. Some trials showed power at higher harmonics as well. B) Δ*d*_10_ and C) stress correlated with a phase shift. In this case muscle 178 had a Pearson correlation coefficient, *r* = −.30 (*p* = .002) and muscle 179 had *r* = −.31 (*p* = .0009). D) Incremental work calculated as σ_*a*-*p*_(*t*)Δ ∊ (*t*) = *W*_*inc*_ (*t*), where σ_*a*-*p*_(*t*) represents active − passive stress. For the 8 Hz conditions, n=5 for muscle 178, and n=7 for muscle 179. For the 11 Hz conditions, n=4 for muscle 178, and n=6 for muscle 179.

### Lattice spacing dynamics correlate to changes in work

Given the lattice spacing difference between muscle 178 and 179, we next tested whether these changes correlated to the timing of stress differences in the two muscle’s dynamic behavior. We could not exactly prepare the muscles in the same ways as in the experiments from ***Ahn et al. (2006)*** where the muscle was left *in situ* in the limb and the motor neuron directly stimulated. To restrict x-ray imaging to a single muscle, work loop preparations in the beamline required isolating the muscles from the cockroach leg and directly stimulating them with silver wire electrodes (***Sponberg et al., 2011a***). When extracellularly stimulating, muscle force rise times are sooner (estimated at 8 ms) because of the lack of transmission and synaptic delays and fall off sooner, likely because all sarcomeres are simultaneously activated (***Sponberg et al., 2011a***). Consequently, under identical 8 Hz running conditions, force develops sooner in our muscle preparations than in the neural stimulation, *in situ* work loops of ***Ahn et al. (2006)***. As a result, under extracellular stimulation both muscles 178 and 179 produce small but significant positive work and more negative work (Table 1). In prior experiments, faster locomotor trials at 11 Hz were observed and implemented in work loops (***Sponberg et al., 2011a***). In muscle 137, the midleg equivalent of 179 these 11 Hz conditions with extracellular stimulation gave more similar performance to the ***Ahn et al. (2006)*** and ***Full et al. (1998)*** conditions. The faster frequency reduced stride period correspondingly. To compare with these conditions, we repeated all of our trials under 11 Hz work loops. In this case, we found results more consistent with previous work loops. Muscle 178 produced positive work statistically indistinguishable from the 8 Hz condition (*p* = .56, *t*-test), but muscle 179 produced significantly less (*p* = .017, *t*-test) and both muscles produced even more negative work than in the 8 Hz conditions (*p* = .07 and *p* = .002, *t*-test, for muscles 178 and 179, respectively). The differences between the two muscles that we observed are not as dramatic as those from the *in situ* work loops, likely because of the preparation differences. However, negative work also has large variation (50-75%) from experiment to experiment in both our experiments and previous studies at these conditions (***Ahn et al., 2006***; ***Sponberg et al., 2011a***).

**Table 1.**
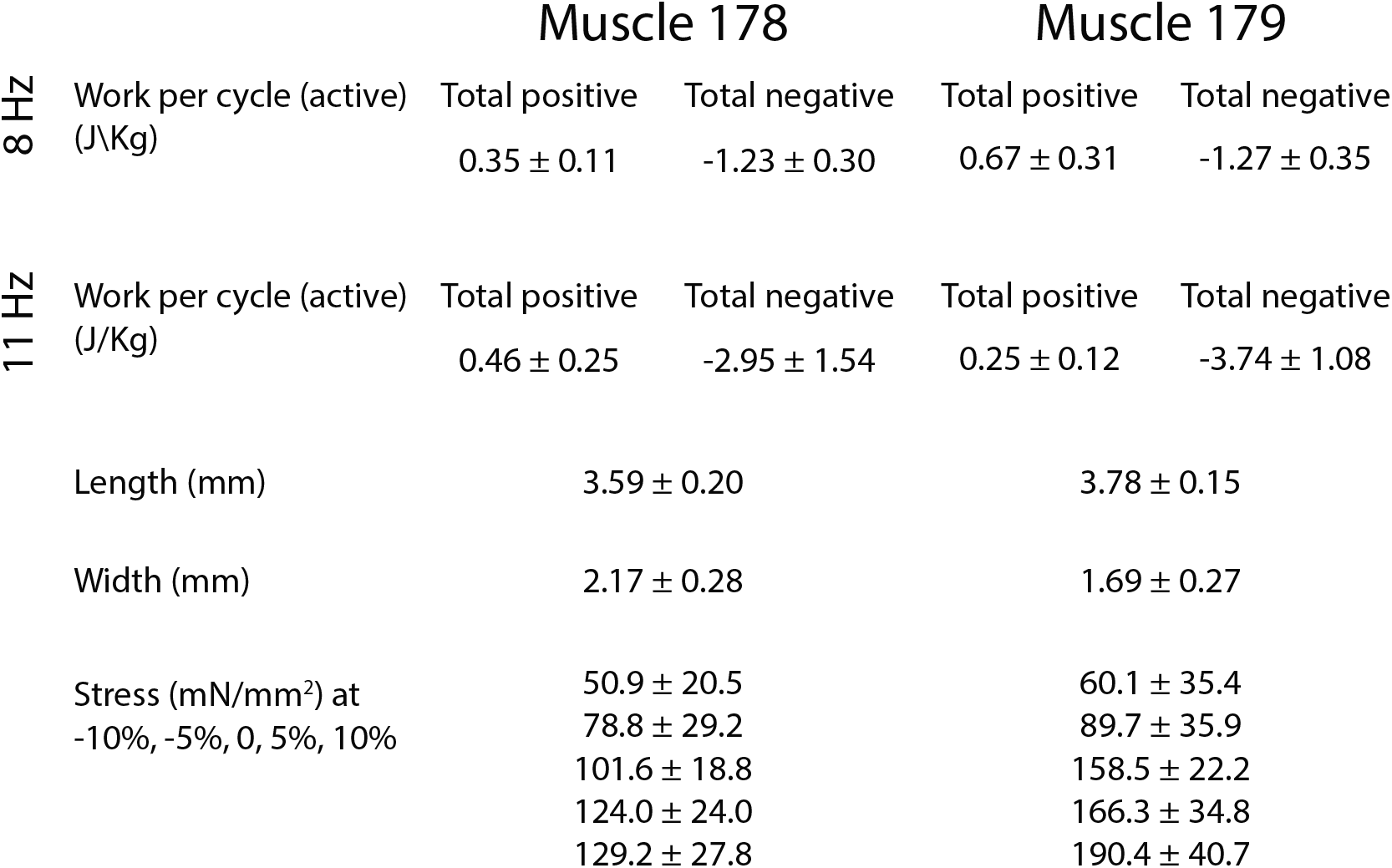
All values are means ±95% confidence intervals of the mean. For the 8 Hz conditions, n=6 for muscle 178, and n=7 for muscle 179. For the 11 Hz conditions, n=4 for muscle 178, and n=9 for muscle 179.

Given the variation, we considered the correlations between lattice spacing and stress in every individual trial from both the 8 Hz and 11 Hz work loops. We paired active and passive work loop conditions for each individual and first tested if the difference in lattice spacing due to activation, Δ*d*_10_ = *d*_10, *active*_ − *d*_10, *passive*_, was periodic at the underlying work loop frequency (8 or 11 Hz). We detrended and took the Fourier transform of Δ*d*_10_ for each individual experiment. We calculated the phase and power spectrum for each. In all cases there was significant power in Δ*d*_10_ at the work loop frequency, although in some trials there was also signal at the harmonics. Under the 8 Hz work loop conditions, from the Fourier series, we determined the average phase shift between stress (active - passive) and Δ*d*_10_ to be −12.3 ± 17.3 ms for muscle 178 and −22.3 ± 14.1 ms for muscle 179 (mean ± 95% CI of the mean) with a negative phase shift indicating stress precedes Δ*d*_10_. A phase shift makes sense given that the lattice spacing change on top of that due to passive axial strain, likely arises from the myosin crossbridges producing radial force. The phase was not significantly different between the two muscles. Under the 11 Hz conditions, we found a significant (*p* = .02, *t*-test) difference between the average phase shift between stress and *d*_10_ to be −16.3 ± 34.5 ms for muscle 178 and 25.6 ± 11.7 ms for muscle 179.

To align the stress, we shifted the Δ*d*_10_ to the frame closest to the average phase measured in the power spectra. In all 8 Hz and 11 Hz trials, changes in lattice spacing from passive to active work loop conditions correlated with stress. Figure 5 shows a representative time series of Δ*d*_10_, stress, and incremental work for muscle 178 and 179 at 8 Hz. Peak Δ*d*_10_ under the 8 Hz conditions in muscle 178 was larger than in 179. Muscle 178’s increased lattice spacing change corresponded with a plateau in stress, compared to the stress in muscle 179 which rises rapidly and falls.

### Lattice spacing dynamics depend on strain offset

Under perturbed conditions during locomotion these muscles can undergo many different patterns of strain. We next changed mean strain offset to test if changes in mean strain had a large effect on the lattice spacing dynamics during the work loops. A homologous muscle to 179 has a large functional range, shifting from a brake to a motor under different activation and strain conditions (***Sponberg et al., 2011a***). If lattice spacing covaries with work, we would expect large variation in lattice spacing dynamics under different strain conditions.

The difference in lattice spacing dynamics between the two muscles was present at every strain condition. The peak-to-peak amplitude of *d*_10_ in muscle 178 always increased during activated work loops compared to passive conditions (figure 7). This change was larger than the Δ*d*_10_ for muscle 179 in every case except at −5%, where *d*_10_ decreased in muscle 179. However, muscle 179 showed a much greater sensitivity to mean strain. In many cases the lattice spacing was actually reduced when the muscle was activated, indicating that myosin activation constrained the radial expansion of the lattice.

## Discussion

A single nanometer difference in the myofilament lattice of two otherwise identical muscles can account for one of the muscles acting like a brake, while the other produces more positive mechanical work. Before activation, *d*_10_ in muscle 178 has a smaller lattice spacing than muscle 179 by approximately 1 nm at 10% strain, which is where activation occurs *in vivo* (figure 8). Simply showing that there is a passive lattice spacing difference is insufficient to explain the two muscles’ different work production because under steady state (isometric and isotonic) conditions, these two muscles produce the same force. However, the 1 nm lattice spacing difference disappears during isometric twitches, consistent with the identical steady state macroscopic properties. Once stimulated identically and held isometrically as in ***Ahn et al. (2006)***, 178 pushes the lattice further apart, whereas muscle 179 is already at its steady state lattice spacing (see figure 8, green dotted lines).

**Figure 8.**
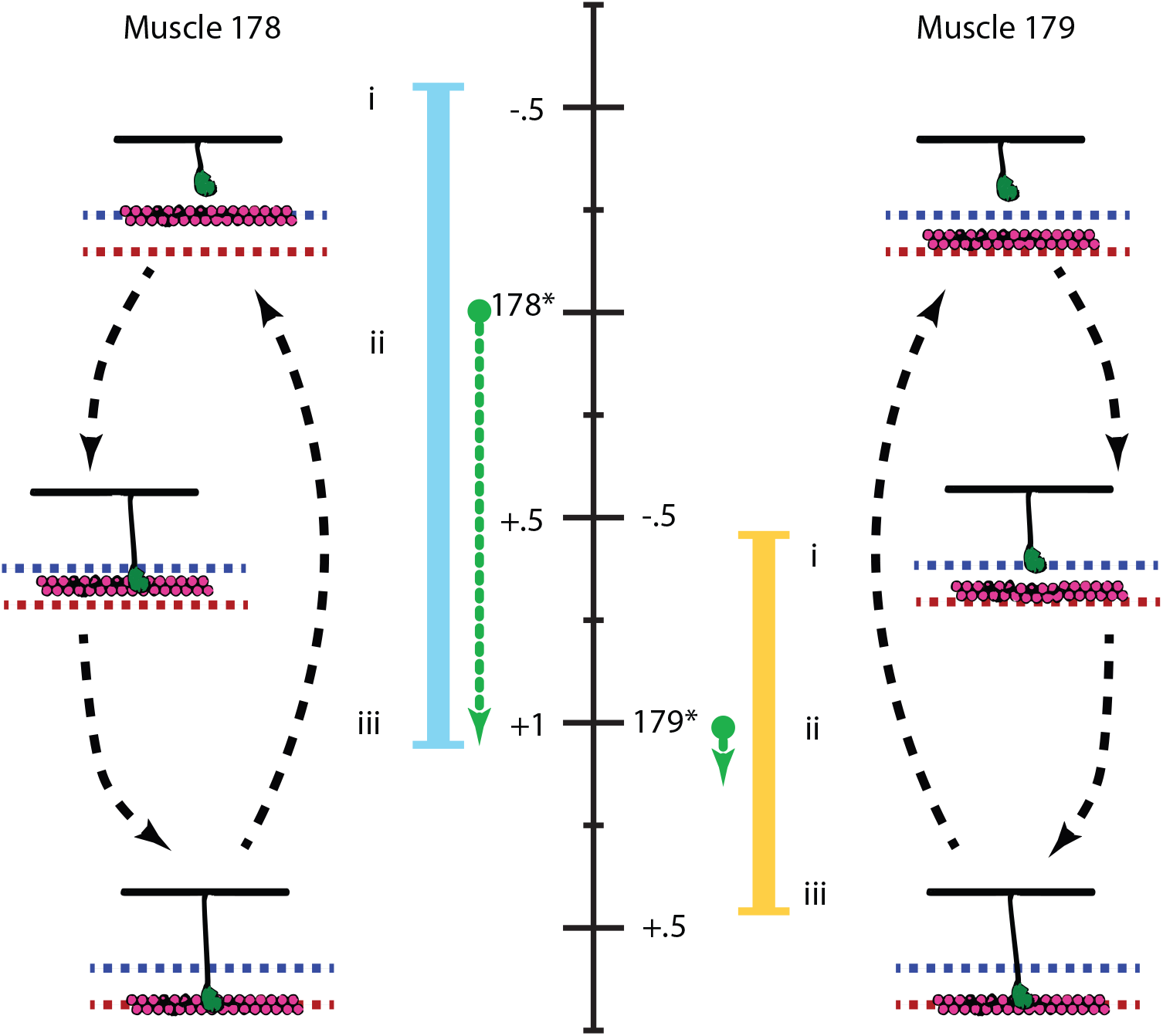
Black dashed arrows illustrate the crossbridge and lattice spacing states during activation. The black scale (in nm) shows the range of lattice spacing change for both muscles, with their values at rest indicated by (*), and centered around muscle 178’s rest value. Under isometric conditions, the lattice spacing in muscle 178 increases while muscle 179’s does not, leaving them at the same lattice spacing at peak activation (green dashed lines). During passive, unactivated work loops, lattice spacing changes due to axial strain (Figure 4). We subtracted that passive cycling off to show the difference in lattice spacing due solely to activation of muscle during workloops, Δ*d*_10_ (solid blue and yellow lines). The timing of activation was near the start of shortening (*i*). Before this time the muscles are unactivated, offset by 1 nm, and muscle 178 has a 1 nm tighter lattice spacing denoted by the dashed blue line. During early shortening (*ii*) muscle 178 produces more positive work, likely because it is in a more favorable position for myosin heads to bind, and undergoes a larger transient in lattice spacing change. By the end of shortening (*iii*) and into lengthening, the myosin heads have bound and pushed the thin filaments (pink) out to the steady state value (red dashed line). This expansion is greater in muscle 178. So for both steady state (peak activation during isometric conditions) and dynamic (whole work loop), muscle 179’s lattice spacing is greater, but more constrained, while muscle 178’s is smaller but undergoes a greater range of lattice spacing change. These differences in lattice spacing can account for the similarities in their steady state macro-scale properties (dashed green lines end at the same point) as well as the difference in their mechanical work production (blue and yellow lines are different).

The consequence of this lattice spacing difference that disappears under active isometric conditions is that muscle 178 undergoes a 0.82 nm larger change in lattice spacing during periodic contractions compared to muscle 179 (figure 8). Since the amount of force that is generated axially is dependent on the lattice spacing, as is the crossbridge binding probability ***Schoenberg (1980)***; ***Williams et al. (2010)***, this increased change in lattice spacing can have functional consequences. Even though constraints on doing work loops within the x-ray beamline required different methods of stimulation and muscle preparation, changes in lattice spacing correlate with changes in work production in both muscles 178 and 179. The increased transient change in 178’s *d*_10_ after activation corresponds to the plateau in stress development during this portion of the contraction cycle. We cannot currently manipulate lattice spacing within intact muscle independent of cross bridge activity to causally connect to muscle function. However, our results can explain both the dynamic differences and the steady state similarities of these two cockroach muscles.

The coupling of lattice spacing and muscle stress production is complicated because the coupling of lattice to work happens across the hierarchy of muscle organization, and it is not understood how one length scale couples to another. Spatially explicit models have shown that lattice spacing can affect force, but these models cannot yet predict work under dynamic conditions for a full 3-D lattice (***Williams et al., 2010***; ***Tanner et al., 2007***). Other detailed half-sarcomere models can capture work differences but cannot yet explicitly incorporate myofilament lattice differences (e.g. ***Campbell et al. (2011b,a)***. While we cannot yet predict the differences in work, our results link nanometer scale structural differences with functional differences relevant for locomotion.

### Packing structure cannot account for the differences in these two muscles

Because no statistically significant difference was found in the measurements we took of 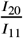 for the two muscles, we determined the two muscles to have the same ratio and arrangement of myosin to actin filaments. Since the muscles are both femoral extensors acting at the same joint, it might seem natural to assume from the beginning that they have the same packing structure. However, even though *B. discoidalis* is flightless, electron micrographs have shown that the largest of the femoral extensors in the middle leg which is in between the homologs of these two muscles actually has fight muscle packing arrangement (***Jahromi and Atwood, 1969***). This is presumably because this muscle is bifunctional and also actuates the wings (***Carbonell, 1947***). It has also been shown that wing actuation muscles in the beetle *Mecynorrhina torquata* which act as steering muscles have limb muscle architecture (***Shimomura et al., 2016***). So it is not always possible to assume a given packing geometry based only on muscle function.

Although the packing pattern of these two cockroach muscles does not explain their work loop differences, it is still an open question how different packing structures might affect muscle function and energetic versatility. Structure indeed does seem to be related to function. In vertebrate muscle (human gastrocnemius (***Widrick et al., 2001***), rabbit psoas (***Hawkins and Bennett, 1995***), frog sartorius (***Luther and Squire, 2014***), all seen by electron microscopy, and others (***Millman, 1998***; ***Squire et al., 2005***)) actin is arranged such that one actin is located equidistant from 3 myosin, which makes a 1:2 myosin:actin ratio per unit cell. Invertebrate muscle actin packing can vary greatly, with even adjacent muscles in the same animal having different actin arrangement. Flight muscle (*drosophila* (***Irving, 2006***), *Lethocerus cordofanus* (***Miller and Tregear, 1970***)), for example has one actin located equidistant between every 2 myosin, which makes a 1:3 myosin:actin ratio per unit cell, whereas invertebrate limb muscle (crab leg muscle (***Yagi and Matsubara, 1977***), crayfish leg (***April et al., 1971***)) has 12 actin filaments surrounding each myosin, which makes which makes a 1:6 myosin:actin ratio per unit cell. Different packing structures will have different actin-myosin spacing even if *d*_10_ is the same between muscles since the geometry of actin relative to myosin has changed but myosin geometry has not (***Millman, 1998***). Different ratios will also affect the availability of actin binding sites for myosin heads. The broad interspecific correlation with muscle locomotor type suggests that packing structure may still be an important determinant of work, just not in the two cockroach muscles considered here.

### Structural differences at the micro-scale can explain functional differences at the macro-scale

It is perhaps surprising that a 1 nm spacing difference could have such a dramatic functional consequence. Even when we consider the change relative to the absolute lattice spacing of ≈50 nm, it is only a 2% difference (figure 3). However small differences in myofilament configuration can have dramatic effects because of the sensitivity of myosin’s spatial orientation relative to its binding site on the thin filament. Crossbridge kinetics depend on lattice spacing and vice versa (***Schoenberg, 1980***; ***Adhikari et al., 2004***; ***Tanner et al., 2007***; ***Williams et al., 2013***). By undergoing a larger range of lattice spacing during a typical contraction, muscle 178’s crossbridge kinetics will change more than 179’s crossbridge kinetics.

It is not unprecedented for lattice spacing changes to have multiscale physiological consequences. Temperature has been shown to affect crossbridge activity enough to change *d*_10_ by as much as 1 nm in hawk moth flight muscle (***George et al., 2013***). In that case the temperature difference also corresponds to a functional difference where the cooler superficial part of the muscle acts like a spring while the warmer interior does net positive work. In the cockroach muscles there is unlikely to be any temperature difference because both muscles are small and superficial. While the origin of the lattice spacing differences in these muscles is unknown, it is reasonable that a 1 nm difference in lattice spacing could influence crossbridge activity enough to make a sizable change in work output.

The importance of a 1 nm difference in lattice spacing reflects the more general feature of muscle’s multiscale nature. Multiscale effects manifest when there is coupling between different length scales and when physiological properties arise which are not predicted by the behavior of other length scales. As myosin crossbridges form, lattice spacing can change due to the radial forces generated, aiding or impeding further crossbridge attachment (***Williams et al., 2010***). Also, crossbridge formation strains myosin thick filaments axially, which can influence myosin cooperativity (***Tanner et al., 2007***). This means crossbridges (10’s of nanometer scale) influence and are influenced by the arrangement and strain on the whole sarcomere (micron scale). The deformation of the sarcomere is also a product of strain imposed on the whole muscle fiber (100s of microns), which introduces coupling between whole muscle dynamics and crossbridge kinetics. As an example of physiological effects emerging at different scales, we generally cannot yet predict mechanical work from steady-state physiological properties, especially during perturbed conditions.

### How might different time courses of lattice spacing arise?

Lattice spacing changes are variable across different muscles. In frog muscles the lattice is isovolumetric as rest (***Matsubara and Elliot, 1972***) and in active indirect flight muscle lattice change is minimal (***Irving and Maughan, 2000***). However, our results show that under some strain conditions (see Figure 6, 0 and +5% strain offset in muscle 178) even passive muscle is not strictly isovolumetric, and that the lattice spacing increase after activation can make muscles more isovolumetric. This indicates that individual muscles might have different dependencies on length change as well as activation, as we see in Figure 7.

**Figure 6.**
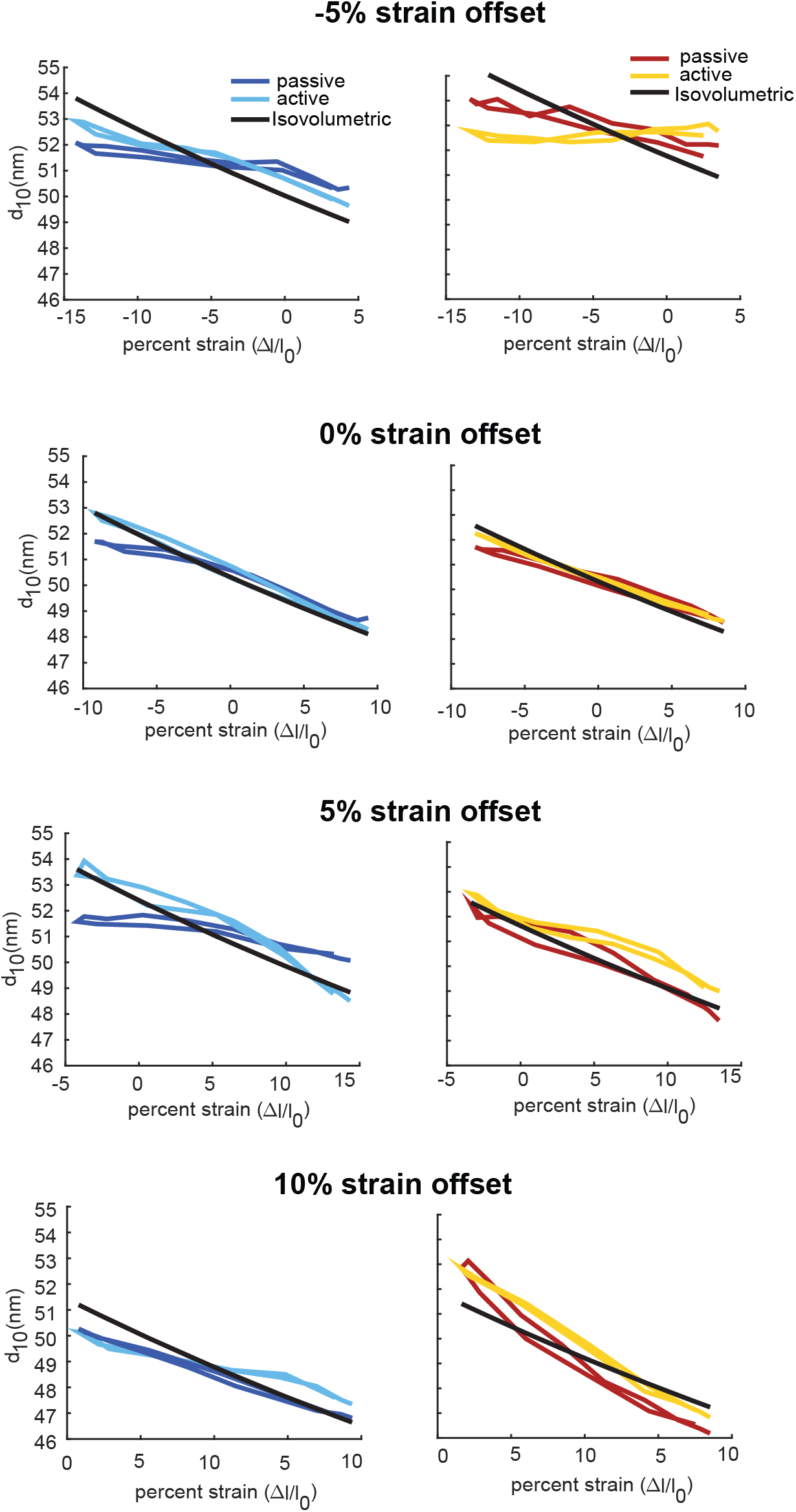
Mean lattice loops (strain vs. *d*_10_) at strain offsets of −5%, +0%, +5%, +10% (top to bottom) for muscles 178 and 179 (left and right). The lattice spacing change in passive conditions is due solely to the axial strain of the myofilament lattice during compression and tension. Under activated conditions the spacing patterns change due to the action of active myosin binding. Sample size n for strain conditions (−5,0,5,10) was: passive muscle 178, n=5 for all strains; active muscle 178, n=(5,6,5,5); passive and active muscle 179, n=(5,8,8,5).

**Figure 7.**
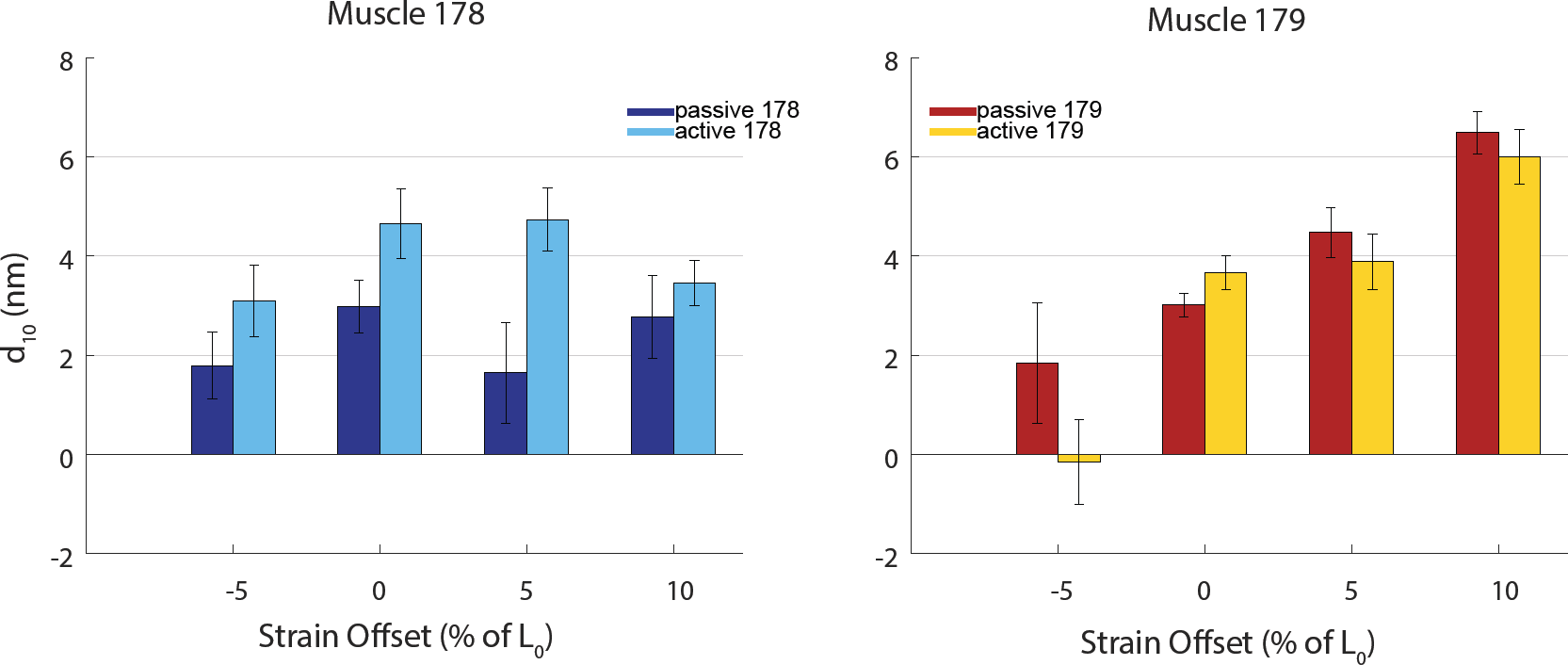
Mean change in lattice spacing from start of shortening to end of shortening with 95% confidence of the mean for muscles 178 (left) and 179 (right) during passive and active work loops. We found that strain greatly affected lattice spacing for muscle 179 (*p <* .001), but not for muscle 178 (*p* = .43). In contrast, we found activation greatly affected muscle 178 (*p* = .007) but did not significantly affect muscle 179 (*p* = .24). Statistics were calculated by 2-factor ANOVA. See Figure 6 for sample sizes.

Many experiments have shown that the relationship between sarcomere length and lattice spacing may be regulated by titin (***Fuchs and Martyn, 2005***). For example, by enzymatically lowering the passive tension of titin in mice, it was seen that lattice spacing increased and pCa sensitivity decreased, implying there exists a strong radial component of titin force which in2uences actin-myosin interaction possibly by regulating the lattice structure (***Cazorla et al., 2001***). Bovine left ventricles and left aortas express higher and lower titin stiffness, respectively. Ca^2+^ sensitivity with sarcomere length is much stronger in the ventricle with stiffer titin, and this is coupled with smaller lattice spacing, as seen with x-ray diffraction (***Fukuda et al., 2003***). In the muscles in our study, lattice spacing differences might be explained by differences in projectin or sallimus, the titin-like proteins found in insects ***Yuan et al. (2015)***. Muscle 179 having stiffer titin-like proteins would be consistent with these previous results because in that muscle the myofilament lattice spacing has a greater dependence on length (Figure 7).

The offset in filament spacing between the two muscles could also arise from differences in Z disk proteins, like α-actinin, which cross-link actin (***Hooper and Thuma, 2005***). While this could account for the passive offset it is less clear how such structural differences in the anchoring of actin alone could also explain why the *d*_10_ difference between the two muscle disappears under steady state activation.

### Structural elements of the actin-myosin lattice have implications for understanding control

In addition to similar muscles producing different amounts of mechanical work under comparable conditions, the same muscle can also have a great deal of functional variation. How lattice spacing interplays with macroscopic force production might contribute to the how a muscle changes function under perturbed conditions. The way a muscle’s lattice spacing changes during periodic contractions at different strains give clues to how muscles can achieve such versatile functions. Comparing the lattice loops of passive 178 and 179 in Figure 6, muscle 179’s lattice spacing has a more sensitive dependence on strain, and a smaller dependence on activation compared to muscle 178 (Figure 7). On 2at terrain while running this muscle’s *in vivo* function is to act as a brake. However when perturbed, it can perform large amounts of positive work which can affect center of mass behavior of the whole insect. In muscle 137, the mid-limb analogue of muscle 179, a large change in function can arise from small changes in strain and phase of activation which arise from either neural or mechanical feedback (***Sponberg et al., 2011b***,a). By having lattice spacings with different dependencies on muscle length and activation, different muscles may be able achieve large functional variation such as muscle 137, or be robust in their function even as activation changes.

## Conclusion

A 1 nm difference in the spacing of the myofilament lattice is the first feature that can account for the functional difference of two nearly identical leg muscles in the cockroach. Nanometer differences in lattice spacing not only influence myosin binding, but may explain categorical shifts in muscle function that have effects at the scale of locomotion. A single nanometer change in spacing can have this profound effect because of the multiscale coupling from the molecular lattice to the tissue. Simultaneous time resolved x-ray diffraction and physiological mechanism are starting to link biophysical differences in muscle structure to macroscopic function even under dynamic conditions.

## Methods and Materials

### Animals

*Blaberus discoidalis* were maintained in a colony at Georgia Tech under a 12:12 light dark cycle and provided food *ad libitum*. Muscles 178 and 179 are located on the mediodorsal and medioventral sides of the coxa respectively (***Ahn et al., 2006***). After removing the whole hind-limb, the leg was pinned such that the femur formed a 90° angle with the axis of contraction for 178 and 179 with either dorsal or ventral side facing up. After removing enough exoskeleton to view the muscle of interest, its rest length (RL) was measured from a characteristic colored spot on the apodeme to the anterior side of the coxa where the muscle originates (***Full et al., 1998***). We also measured the width of the muscle at mid-length. Once dissected from the coxa, the muscle was mounted between a dual-mode muscle lever (model 305C, Aurora Scientific, Aurora, Canada) and a rigid hook, and length was set to 104.4% RL for muscle 178 and 105% for muscle 179 - this defined the operating length (OL) of the muscle, or the mean length during *in vivo* running (***Ahn et al., 2006***; ***Ahn and Full, 2002***). Silver wire electrode leads were placed at opposite ends of the muscle for extra-cellular activation as in (***Sponberg et al., 2011a***).

### Time Resolved x-ray Diffraction

Small angle X-ray diffraction was done using the Biophysics Collaborative Access team (BioCat) small angle diffraction instrument on Beamline 18ID at the Advanced Photon Source (APS), Argonne National Laboratory. The beam dimensions at the focus were 60 × 150 *μm*, vertically and horizontally respectively with a wavelength of .103 nm (12 keV). Initial beam intensity is 10^13^ photons/s, which we attenuated with 12 sheets of 20 *μm* thick aluminum, about a 65% reduction. For all cases, diffraction images were recorded on a Pilatus 3 1M pixel array detector (Dectris Inc) with an exposure time of 4 ms with a 4 ms period between images during which a fast shutter was closed to reduce radiation damage.

### Experimental Protocol

After being extracted and mounted, muscles were placed in the beam-line and activated with a twitch consisting of 3 spikes separated by 10 ms, with the first occurring at *t* = 0 ms. Diffraction images we collected starting from *t* = −25 ms and ending at *t* = 175 ms. One twitch was done at strain offsets of −10, −5, 0, +5, +10% OL each for both muscles. We estimated cross-sectional area from the diameter of the muscle assuming a cylindrical shape, and used this to calculate stress. From this we obtained the lattice spacing *d*_10_ during the whole twitch.

Next, we did work loops under several conditions. First, strain amplitude (peak to peak) was 18.5% of OL for muscle 178 and 16.4% of OL for muscle 179, with different strain amplitudes accounting for different absolute lengths. The driving frequency was 8 Hz, with activation consisting of 3 spikes at 6 volts at 100 Hz, at a phase of activation of .08%, with 0 defined as the start of shortening. These are the *in vivo* conditions of these muscles during running (***Full et al., 1998***; ***Ahn et al., 2006***), except with the muscle isolated and extracellularly stimulated. We also did work loops at 11 Hz with the same activation, which matches the conditions from ***Sponberg et al. (2011a)*** including the same method of stimulation. We then performed the same work loop but with strain offsets of −10, −5, 0, +5, +10 percent OL, and did passive work loops for every active work loop. Each work loop trial consisted of 8 cycles, and we discarded the first cycle. Muscle stress was calculated using the average mass values from (***Ahn et al., 2006***) and the measured resting lengths because these measurements produced less variation than attempts to measure mass following x-ray experiments. During our limited beam time we had 17 total samples.

### Analysis

The most prominent peaks in the muscle diffraction patterns are the (1,0), (1,1), (2,0) peaks, all of which correspond to planes in the muscle crystal lattice (see Figure 1 C and E). Since the intensity is related to the mass which lies along the associated plane, we can use the (1,1) and (2,0) peaks to determine the arrangement of actin in the lattice. If more mass is located along the (1,1) plane, as in vertebrate muscle, the (1,1) peak will be much brighter than the (2,0) peak, and 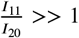. In invertebrate flight muscle, more mass is aligned with the (2,0), which will mean the (2,0) peak is brighter than the (1,1): 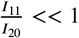(***Irving, 2006***). The spacing between two peaks gives the spacing between the corresponding planes in the lattice via Bragg’s Law, so we can use the (1,0) peaks to determine the lattice spacing *d*_10_.

X-ray diffraction patterns were analyzed by automated software (***Williams et al., 2015***), a subset of which was verified by hand fitting with *fityk*, a curve fitting program (***Wojdyr, 2010***). Individual frames for which the automated software failed to resolve peaks were discarded. Trials with frames that consistently failed during multiple cycles to resolve peaks were discarded totally.

## Acknowledgments

The authors thank George Steven Chandler and Chidinma Chukwueke for help in data collection and Sage Maligen, Tom Libby, and Tom Daniel for helpful discussions. This work was supported by grant W911NF-14-1-0396 from the Army Research Office to SNS, TCI, Tom Daniel and Anette Hosoi. This research used resources of the Advanced Photon Source, a U.S. Department of Energy (DOE) Office of Science User Facility operated for the DOE Office of Science by Argonne National Laboratory under Contract No. DE-AC02-06CH11357. The work was supported by GUP beamtime awards 47291 and 52213. Use of the Pilatus 3 1M detector was provided by grant 1S10OD018090-01 from NIGMS. This project was also supported by grant 9 P41 GM103622 from the National Institute of General Medical Sciences of the National Institutes of Health. The content is solely the responsibility of the authors and does not necessarily reflect the official views of the National Institute of General Medical Sciences or the National Institutes of Health.

## References

Adhikari BB, Regnier M, Rivera A J, Kreutziger KL, Martyn DA. Cardiac Length Dependence of Force and Force Redevelopment Kinetics with Altered Cross-Bridge Cycling. Biophysical Journal. 2004 Sep; 87:1784–1794.

Ahn AN. How Muscles Function-the Work Loop Technique. Journal of Experimental Biology. 2012; 215(7):1051–1052. doi: 10.1242/jeb.062752.

Ahn AN, Full RJ. A motor and a brake: two leg extensor muscles acting at the same joint manage energy differently in a running insect. Journal Of Experimental Biology. 2002; 205(3):379–389.

Ahn AN, Meijer K, Full RJ. *In situ* muscle power differs without varying *in vitro* mechanical properties in two insect leg muscles innervated by the same motor neuron. Journal Of Experimental Biology. 2006; 209(17):3370–3382.

April EW, Brandt PW, Elliott GF. The myofilament lattice: Studies on isolated fibers. The Journal of Cell Biology. 1971; 51(1):72–82. doi: 10.1083/jcb.51.1.72.

Bagni MA, Cecchi G, GriZths PJ, Maeda Y, Rapp G, Ashley CC. Lattice spacing changes accompanying isometric tension development in intact single muscle fibers. Biophys J. 1994 Nov; 67(5):1965–1975.

Becht G, Dresden D. Physiology of the Locomotory Muscles in the Cockroach. Nature. 1956 jan; 177(4514):836–837.

Becht G, Hoyle G, Usherwood PNR. Neuromuscular transmission in the coxal muscles of the cockroach. Journal of Insect Physiology. 1960 aug; 4(3):191–201. doi: 10.1016/0022-1910(60)90026-3.

Campbell SG, Hatfield C, Campbell KS. A model with Heterogeneous Half-Sarcomeres Exhibits Residual Force Enhancement After Active Stretch. Biophysical Journal. 2011; 100(3):12a. http://dx.doi.org/10.1016/j.bpj.2010.12.277, doi: 10.1016/j.bpj.2010.12.277.

Campbell SG, Hatfield PC, Campbell KS. A Mathematical Model of Muscle Containing Heterogeneous Half-Sarcomeres Exhibits Residual Force Enhancement. Plos Computational Biology. 2011 sep; 7(9):e1002156. http://dx.plos.org/10.1371/journal.pcbi.1002156.g008.

Carbonell CS. The Thoracic Muscles of the Cockroach Periplaneta Americana (L.). Smithsonian Miscellaneous Collections. 1947 jan; 107(2):1–23.

Cazorla O, Wu Y, Irving T, Granzier H. Titin-Based Modulation of Calcium Sensitivity of Active Tension in Mouse Skinned Cardiac Myocytes Materials and Methods Preparations and Solutions. Circulation Research. 2001; 88(10):1028–1035. doi: 10.1161/hh1001.090876.

Cecchi G, Bagni M, GriZths P, Ashley C, Maeda Y. Detection of radial crossbridge force by lattice spacing changes in intact single muscle fibers. Science. 1990 Dec; 250:1409–1411.

Dickinson MH, Farley CT, Full RJ, Koehl MA, Kram R, Lehman S. How animals move: an integrative view. Science. 2000 Apr; 288(5463):100–106.

Fuchs F, Martyn DA. Length-dependent Ca2+ activation in cardiac muscle: some remaining questions. Journal of Muscle Research and Cell Motility. 2005 Apr; 26:199–212.

Fuchs F, Wang YP. Sarcomere Length Versus Interfilament Spacing as Determinants of Cardiac myofilament Ca2+Sensitivity and Ca2+Binding. Journal of Molecular and Cellular Cardiology. 1996; 28(7):1375–1383. doi: https://doi.org/10.1006/jmcc.1996.0129.

Fukuda N, Wu Y, Farman G, Irving TC, Granzier H. Titin Isoform Variance and Length Dependence of Activation in Skinned Bovine Cardiac Muscle. The Journal of Physiology. 2003; 553(1):147–154. doi: 10.1113/jphys-iol.2003.049759.

Full RJ, Stokes DR, Ahn AN, Josephson RK. Energy absorption during running by leg muscles in a cockroach. Journal Of Experimental Biology. 1998 Apr; 201 (Pt 7):997–1012.

George NT, Irving TC, Williams CD, Daniel TL. The cross-bridge spring: can cool muscles store elastic energy? Science. 2013 Jun; 340(6137):1217–1220.

Gordon AM, Huxley AF, Julian FJ. The variation in isometric tension with sarcomere length in vertebrate muscle 1bres. The Journal of Physiology. 1966; 184(1):170–192. doi: 10.1113/jphysiol.1966.sp007909.

Hawkins CJ, Bennett PM. Evaluation of freeze substitution in rabbit skeletal muscle. Comparison of electron microscopy to X-ray diffraction. Journal of Muscle Research & Cell Motility. 1995; 16:303–318.

Hooper SL, Thuma JB. Invertebrate Muscles: Muscle specific Genes and Proteins. Physiological reviews. 2005 jul; 85(3):1001–1060.

Huxley AF, Simmons RM. Proposed Mechanism of Force Generation in Striated Muscle. Nature. 1971 Oct; 233:533–538.

Irving TC, Maughan DW. In Vivo X-Ray Diffraction of Indirect Flight Muscle from Drosophila melanogaster. Biophysics Journal. 2000 May; 78:2511–2515.

Irving TC. X-ray diffraction of indirect flight muscle from Drosophila in vivo. In: Vigoreaux J, editor. Nature’s Versatile Engine: Insect Flight Muscle Inside and Out Landes Bioscience; 2006.p. 197–211. doi: 10.1007/0-387-31213-7.

Iwamoto H. Synchrotron radiation x-ray diffraction techniques applied to insect 2ight muscle. International Journal of Molecular Sciences. 2018; 19(6). doi: 10.3390/ijms19061748.

Jahromi SS, Atwood HL. Structural Features of Muscle Fibers in the Cockroach Leg. Journal of Insect Physiology. 1969 Jun; 15:2255–2262.

Josephson RK. Mechanical power output from striated-muscle during cyclic contraction. Journal Of Experimental Biology. 1985; 114(JAN):493–512.

Josephson RK. Dissecting muscle power output. Journal Of Experimental Biology. 1999; 202(23):3369–3375.

Luther P, Squire J. The Intriguing Dual Lattices of the Myosin Filaments in Vertebrate Striated Muscles: Evolution and Advantage. Biology. 2014 12; 3:846–865. doi: 10.3390/biology3040846.

Matsubara I, Elliot GF. X-ray diffraction studies on skinned single fibres of frog skeletal muscle. Journal of Molecular Biology. 1972 Dec; 72:957–669.

Maughan DW, Vigoreaux JO. An Integrated View of Insect Flight Muscle: Genes, Motor Molecules, and Motion. Physiology. 1999 jun; 14(3):87–92. doi: 10.1152/physiologyonline.1999.14.3.87.

McCulloch AD. Systems Biophysics: Multiscale Biophysical Modeling of Organ Systems. Biophysical Journal. 2016; 110(5):1023–1027. doi: 10.1016/j.bpj.2016.02.007.

Miller A, Tregear RT. Evidence concerning Crossbridge Attachment during Muscle Contraction.. 1970; 226(5250):1060–1061. doi: 10.1038/2261060a0, exported from https://app.dimensions.ai on 2019/03/16.

Millman B. The Filament Lattice of Striated Muscle. Physiological Reviews. 1998 Apr; 78:359–391.

Pearson KG, Iles JF. Innervation of coxal depressor muscles in the cockroach, Periplaneta americana. Journal of Experimental Biology. 1971 feb; 54(1):215–232.

Roberts TJ, Marsh RL, Weyland PG, Taylor CR. Muscular Force in Running Turkeys: The Economy of Minimizing Work. Science. 1997 Feb; 275:1113–1114.

Schoenberg M. Geometrical factors influencing muscle force development. II. Radial forces. Biophysical Journal. 1980 Apr; 30:69–77.

Shimomura T, Iwamoto H, Vo Doan TT, Ishiwata S, Sato H, Suzuki M. A Beetle Flight Muscle Displays Leg Muscle Microstructure. Biophysical Journal. 2016 09; 111:1295–1303. doi: 10.1016/j.bpj.2016.08.013.

Sponberg S, Libby T, Mullens C, Full R. Shifts in a single muscle’s control potential of body dynamics are determined by mechanical feedback. Philosophical Transactions of the Royal Society B. 2011 May; 366:1606–1620.

Sponberg S, Spence A J, Mullens CH, Full RJ. A single muscle’s multifunctional control potential of body dynamics for postural control and running. Philosophical Transactions of the Royal Society B. 2011 Apr; 366:1592–1605.

Squire JM, Al-khayat HA, Knupp C, Luther PK. Molecular Architecture in Muscle Contractile Assemblies. In: Fibrous Proteins: Muscle and Molecular Motors, vol. 71 of Advances in Protein Chemistry Academic Press; 2005.p. 17–87. doi: https://doi.org/10.1016/S0065-3233(04)71002-5.

Tanner BCW, Daniel TL, Regnier M. Sarcomere lattice geometry influences cooperative myosin binding in muscle. Plos Computational Biology. 2007 jul; 3(7):1195–1211.

Tanner BCW, Farman GP, Irving TC, Maughan DW, Palmer BM, Miller MS. Thick-to-thin filament surface distance modulates cross-bridge kinetics in Drosophila flight muscle. Biophysical Journal. 2012 sep; 103(6):1275–1284. doi: 10.1016/j.bpj.2012.08.014.

Widrick JJ, Romatowski JG, Norenberg KM, Knuth ST, Bain JLW, Riley DA, Trappe SW, Trappe TA, Costill DL, Fitts RH. Functional properties of slow and fast gastrocnemius muscle fibers after a 17-day spaceflight. Journal of Applied Physiology. 2001; 90(6):2203–2211. doi: 10.1152/jappl.2001.90.6.2203.

Williams CD, Regnier M, Daniel TL. Axial and Radial Forces of Crossbridges Depend on Lattice Spacing. PLOS Computational Biology. 2010 Dec; 6(e1001018):1–10.

Williams CD, Salcedo MK, Irving TC, Regnier M, Daniel TL. The length-tension curve in muscle depends on lattice spacing. Proceedings Biological sciences / The Royal Society. 2013 Jul; 280(1766):20130697–20130697.

Williams CD, Balazinska M, Daniel TL. Automated Analysis of Muscle X-ray Diffraction Imaging with MCMC. In: Biomedical Data Management and Graph Online Querying Springer; 2015.p. 126–133.

Wojdyr M. Fityk: a general-purpose peak fitting program. Journal of Applied Crystallography. 2010 Oct; 43:1126–1128.

Yagi N, Matsubara I. The equatorial X-ray diffraction patterns of crustacean striated muscles. Journal of Molecular Biology. 1977; 117(3):797–803. doi: https://doi.org/10.1016/0022-2836(77)90070-5.

Yuan CC, Ma W, Schemmel P, Cheng YS, Liu J, Tsaprailis G, Feldman S, Southgate AA, Irving TC. Elastic proteins in the flight muscle of Manduca sexta. Archives of Biochemistry and Biophysics. 2015; 568:16–27. doi: https://doi.org/10.1016/j.abb.2014.12.033.

